# Interplay of cellular states: Role of delay as control mechanism

**DOI:** 10.1101/630418

**Authors:** Shakti Nath Singh, Athokpam Langlen Chanu, Md. Zubbair Malik, R.K. Brojen Singh

## Abstract

Delay is everywhere, no matter how small or big it is. Experimental evidences show the existence and importance of time delayed reactions specially in biological systems. The role of delay is found to be multifunctional and is seemed to be system dependent. The analytically solved *P*(*X*, *t*) of gene regulatory process shows universal class of Poisson process at stationary condition. However, time delay creates a possible condition to the system to impart correlation in the stochastic process as sub-Poissonian or noise enhancement process which could be important in regulating and controlling the system. The results of simulation of few biological systems (gene regulation, circadian rhythm, and repressilator) using delay stochastic simulation algorithm show the possibilities of delay induced onset of oscillating states, which could be the active states of the systems, where, the system can establish coherence among the system variables, enhance the signal processing, optimize the system activities, etc. On the other hand, delay can also induce switching off of the oscillating states, which may correspond to inactive state or system failure. Further, delay can also create coherent bistable states of the system variables, at which the system can stay longer to make decisions about the fate of the system.

## 1 Introduction

Complex dynamical molecular reaction systems are generally inbuilt with delay. In such homogeneously well-mixed molecular reaction system, incorporation of time delay (*T*) in some of the reactions allows the probability distribution to be at state (*x*, *t*) from an initial state (*x*_*i*_, *t* − *T*) i.e., *P*(*x*, *t*; *x*_*i*_, *t* − *T*) describing the Master equation of the system, to follow non-Markovian process [1]. This time delay plays a vital role in a system to accurately study the dynamical behavior, mechanism to maintain stabilization [2], controlling of signal processing [3] and regulation in the systems [4]. Further, in most biological systems, time delay, which is associated with varied set of reactions having different reaction firing times, can be considered to be an inherent important parameter, and also controls the Master equation describing the dynamics of chemical species. Moreover, time delay, in biological system, is believed to be as a combination of transmission, error detecting, processing and acting times [5]. This biological time delay can be measured experimentally by including maturation and gestation times, voluntary movement reaction times, certain neural reflexes, and the duration to make a protein [6]. There are many biological examples which indicate involvement of delay time. For example, in eukaryotic cells the speed of transcription is approximately twenty nucleotides per second [7], whereas the speed of translation is approximately two codons per second [7]. Human genes are approximately 55,000 nucleotides large [8], and takes 2,750 seconds to transcribe a gene of human [8], however, for translation process, it takes 450 seconds [9]. Other experimental examples show that delay in protein synthesis takes approximately *T* ~ 1 − 2 minutes [10], in bone marrow red blood cells takes *T* ~ 7 days to mature, gestation takes weeks to months, glucose-insulin feedback loop takes of the order of minutes [11], and animals to mature to reproductive age takes months to years [6].

General method for solving chemical Master equation (CME) and delay chemical Master equation (DCME) is the use of generating function method [12], closer form of the CME [13] etc., but for simple cases only. Computationally, modeling each and every details of delay and non-delay chemical reactions involved in the system is computationally intensive. However, delayed chemical reactions can be incorporated as separate events in the CME which could mimic the effects of these minute processes on the overall system dynamics [14]. In these processes which are basically non-consuming reactions [15], a single molecular species may be involved simultaneously in many reaction processes. Hence, the stochastic simulation algorithm (SSA) due to Gillespie, which considers all chemical reactions finish instantly [16–18] and their transients are simple, needs to be modified. To incorporate both delay as well as non-delay reactions in the heterogeneous reactions system, Bratsun et. al. proposed a simulation algorithm to compute both the reaction types systematically [10, 19]. Then to compromise computational cost, Xiaodong Cai proposed a direct method by modifying Bratsun et. al. algorithm [15].

Delay has two contrast roles in regulating deterministic dynamical systems, namely, oscillation death [20] and amplitude death [21] on one hand, and on the other hand, delay induced revival of oscillation [22]. There are reports of delay induced oscillations in stochastic systems, namely, exhibiting oscillations in p53 regulatory network [23?], observation of on and off states of genes in toggle switch [24], emergence of delay induced stochastic oscillations in gene regulation [10] etc. Further, some studies of oscillatory behaviors in Drosophila, Neurospora and other organisms show that these oscillations occur during the transcriptional regulations induced by delay [25–28]. Experiments, then, has proved that these oscillations are caused by induced delay in gene regulation networks [29, 30]. However, the role of delay in inducing oscillations and switching mechanisms in stochastic systems are not fully studied.

In this work, we present a demonstration of delay induced change of cellular phases in some important regulatory models of biological system. We, then, study switching mechanisms/states of oscillations and their relationships with delay time in these models. We provide a detail treatment of stochastic formalism in the models; we study both analytically by incorporating delay parameter, as well as computationally using delay stochastic simulation algorithm. We further present the interplay of delay and noise in the analytical treatment of the systems to highlight the origin of regulating mechanism of delay and noise in the systems. The outline of the paper is as follows. In section 2, we briefly present the delay stochastic master equation formalism and details of delay stochastic simulation algorithm. In section 3, results from analytical work as well as simulation results are presented. We then discuss the results to understand how time delay affects the behavior of the system dynamics and how delay induces the switching mechanisms in stochastic systems. Finally, in section 4, we have concluded by highlighting the important role and effects of time delay in stochastic dynamical systems.

## 2 Materials and Methods

### 2.1 Theoretical framework of delay stochastic master equation

Consider a system of well-stirred *N* molecular species *S* = {*S*_1_,…, *S*_*N*_} which undergo a set of *M* chemical reactions *R*_*i*_: *i* = 1, 2,…, *M*. The state vector of the system at any instant of time *t* is given by *X*(*t*) = [*X*_1_, *X*_2_, *X*_3_,…,*X*_*N*_]^*T*^, where, *X*_*j*_ represents the population of *j*^*th*^ species in the state *X*(*t*). Let *M*_*d*_ and *M*_*n*_ are numbers of delayed and non-delayed reactions such that *M* = *M*_*d*_ + *M*_*n*_. If *τ*_*i*_ = {*τ*_1_, *τ*_2_,…, *τ*_*n*_} are the time delays in the delayed chemical reaction(s) in the probability space with the consideration (*τ*_*i*_ − *τ*_*i*−1_) ~ *f*(*τ*), then the modified master equation which incorporates delay reactions, known as delay stochastic master equation (DSME) [4] is given by,

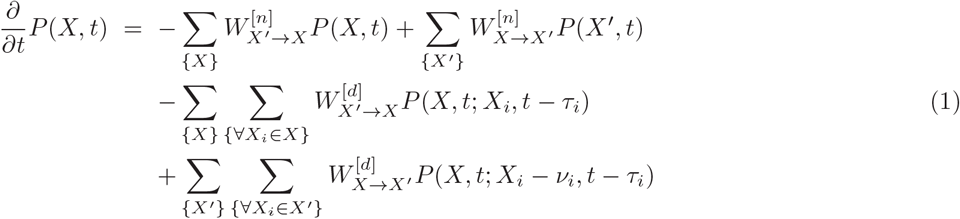

where, *ν*_*i*_ is the stoichiometric or state change ratio. {*W*^[*d*]^} and {*W*^[*n*]^} are delayed and non-delayed transition probabilities. The first two terms in the R.H.S. of the above equation represent non-delay parts of the master equation, and the remaining terms are for delay part. Since, *X*′ = *X* − *ν*_*i*_, one can write,

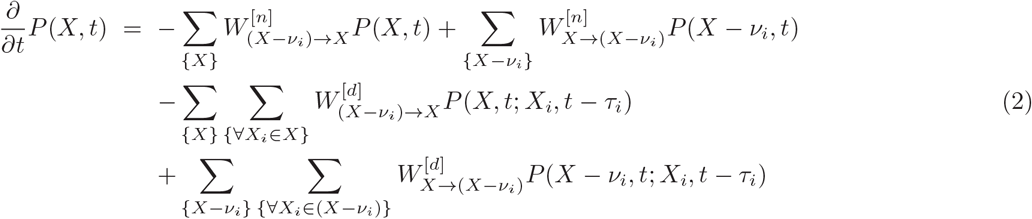

If we consider time delay *τ* to be a distribution of *t*, then equation (2) becomes,

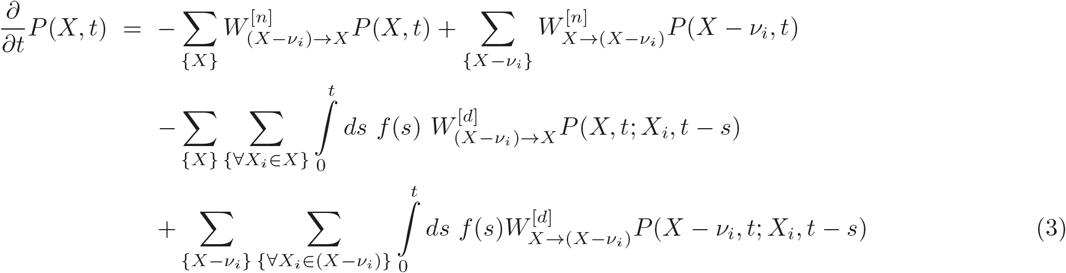

If we assume the coupling between the two states at *t* and (*t* − *s*) to be considerably weak, then the probability distributions at these states could be taken as independent of each other [4], and can be taken as, *P*(*X*, *t*; *X*_*i*_, *t* − *s*) ≃ *P*(*X*, *t*|*X*_*i*_, *t* − *s*)*P*(*X*_*i*_, *t* − *s*). Then using Chapman-Kolmogorov integral equation [31], one can arrive at,

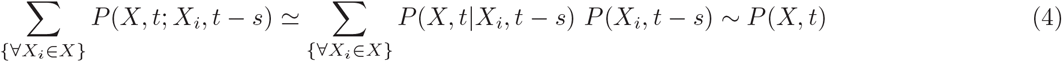

Now, using equation (4), the equation (3) can be written as,

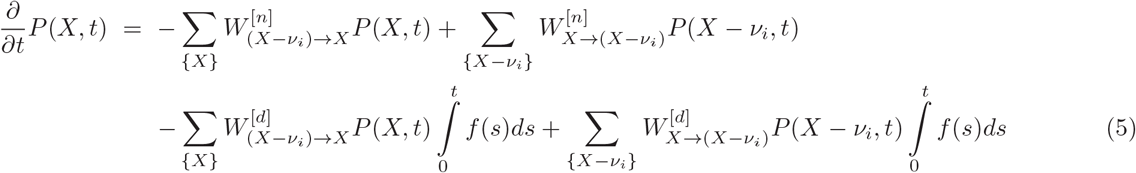

Taking delay as distribution in time *t*, 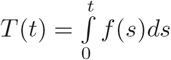, the DSME can be written as,

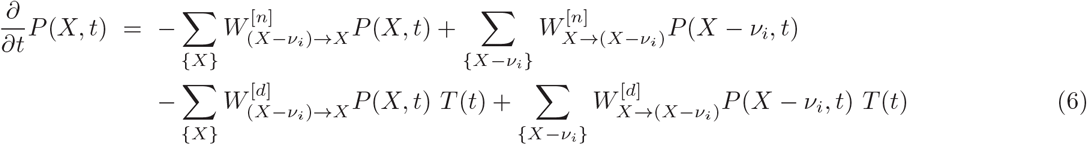

This equation (6), which represents the delay stochastic master equation with the delay distribution function, *T*(*t*), can be solved using generating function technique [12]. Within this framework, one defines a generating function *G*(*s*_1_, *s*_2_,…, *s*_*N*_, *t*) constructed from the probability distribution in equation (6) as given by,

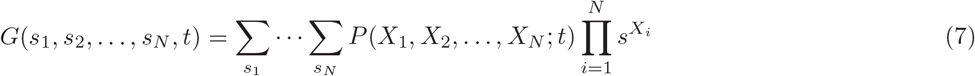

Using equations (6) and (7) one can arrive at the spatio-temporal equation in *G*(*s*_1_, *s*_2_,…, *s*_*N*_, *t*) which can be solved. Substituting the expression for *G*(*s*_1_, *s*_2_,…, *s*_*N*_, *t*) to (7), and using boundary condition *P*(*X*, 0|*X*_0_, 0) = *δ*_*XX*_0__, one can get the expression for *P*(*X*, *t*). The observables, such as ⟨*X*⟩, ⟨*X* ^2^⟩, variance, etc can be calculated using the generating function *G*(*X*_1_, *X*_2_,…, *X*_*N*_).

### 2.2 Delay Stochastic Simulation Algorithm

Complex molecular systems generally consist of broadly two types of reaction viz., delay and non-delay reaction types. Simulations of non-delay reactions, which generally follow Markov process, can be done using stochastic simulation algorithm (SSA) [16–18]. The SSA is based on two important facts during the time evolution of the elementary reactions which is considered to be Markovian process. First, the distribution of reaction times until the next reaction is fired follows 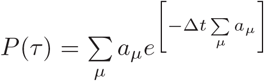, where, *a*_*μ*_ is the propensity function of *μ*^*th*^ reaction given by, *a*_*μ*_ = *h*_*μ*_*c*_*μ*_ [16]. Second, the identification of next reaction (say *j*^*th*^ reaction) to be fired is also a random process, and can be done if the condition 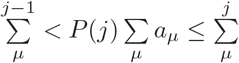 is satisfied. Here, *a*_*μ*_ = *h*_*μ*_*c*_*μ*_, where, *c*_*μ*_ is the stochastic rate constant. These two processes are random processes which are independent events, and hence, satisfy *P*(*τ*, *j*) = *P*(*τ*)*P*(*j*).

The delay reactions generally follow non-Markovian processes [10], and their time evolution trajectories cannot be simulated using SSA. In order to simulate stochastic systems containing delayed reactions, one has to use delayed stochastic simulation algorithm (DSSA) proposed by Bratsun et al. [10] and Barrio et al. [19]. In this simulation process, whenever the delayed reactions occur there will be a change in population of chemical species and the corresponding propensity function will be changed with every time step encountering delay reactions.

The DSSA can be explained as follows. The initialization of the populations of all chemical species is done first. Then the calculation of the propensity function for each reaction is done. Now at each time step, if the reaction encountered is non-delay reaction, then the SSA is used to update the molecular species populations at that time step. If the reaction encountered is delay reaction then we store the finishing time 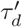 in a list, and we wait till that finishing time to update the state vector for chemical species affected by this delayed reaction [10]. As our simulation time reach to that finishing time of delayed reaction, we update the state vector *X* accordingly, and update the simulation time (*t* + *τ*) with 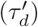 as new time.

There are two types of delayed reactions, namely, non-consuming delayed reactions (in which the reactant of running unfinished reaction can participate in a new chemical reaction) and consuming delayed reactions (in which the reactants of running unfinished reaction can not participate in a new reaction) [19]. The population of reactants of non-consuming delayed reactions will not change until it finishes. Population will change only after the reaction finishes off. In the case of consuming delayed reactions, the population of reactants changes immediately [15]. In delay stochastic simulation algorithm we update state vector accordingly. Because of changes in the state vector of chemical species, we have to recalculate the propensity functions and *τ* for next iteration. Here, *μ* gives the reaction number, which is going to fire in the next time interval [t,t+*τ*). Total number of re-updating time *t* is exactly equal the total number of occurrence of the delayed reactions. It is computationally fast in comparison to other delay stochastic simulation algorithm. But in comparison with direct method for delay Stochastic Simulation, this algorithm generates more random numbers because of delayed reactions [15]. While performing simulation of stochastic models, one faces some challenges. For stochastic simulation we need random number generators as perfect as possible. Simulation result may vary drastically if we use biased random number generator. Some stochastic biological models like polymerase model are very sensitive with initial population of chemical species. Because of this we have found divided by zero error for some model and sometimes it has worked without error. To get more accurate results we need to allow iteration for longer time.

### 2.3 Stochastic Switching Mechanism

The change of state of biological chemical reaction network from one to another state with respect to time due to noise, perturbation, delay, or any biological factor is known as switching mechanism in the biological systems. Due to inherent stochastic nature of the system driven by internal random molecular interaction and environmental fluctuations it allows significant role of noise regulating the system in terms of signal detection and processing, enhancing performance of the system [32], interfering in control mechanism and maintaining robust stabilization of the system [33]. Noise may be internal noise or external noise, which arises in biological networks in the form of random fluctuations [15].

Generally biological systems follow some basic properties like growth, decay and various switchable dynamical states which may correspond to different cellular states. As a result, within a significant time span, they show some dynamical states, such as, various oscillating, stationary, chaotic and transient states and switching on and off among these states. Changes in these states affect the regulatory mechanisms of the biological systems, even may result in genetic switching mechanisms in these biological systems [32]. There are also experimental evidences in the form of bimodal population distributions which show that noise causes the switching from one state to another [32]. Noise may be in many forms. Regulatory mechanisms affecting state changes include positive feedback, double-negative feedback, inhibition /activation, multisite phosphorylation etc [34], and it could be because of the noise, time delay and other parameters which trigger switching behavior in these systems. We intend to analyze this switching behavior in these species variables by using delayed stochastic simulation algorithm.

## 3. Results and Discussions

### 3.1. Delay induced probability distributions of DSME of gene regulation

We present analytical results of gene regulation model (Table 1), where, 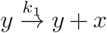 as delay reaction, whereas, 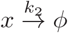 as non-delay reaction in our analytical work, where, *x* and *y* denote polymerase and mRNA species respectively. Now the probability of finding the state *X* at (*t* + Δ*t*) satisfy the following detailed balance condition,

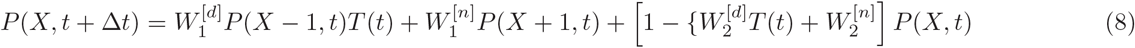

where, the transition probabilities are defined as 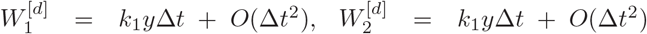, 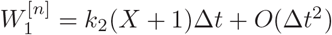 and 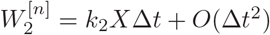 for the two reactions respectively.

#### Theorem 1

*The solution of delay stochastic master equation of gene regulation for the considered model is given by*

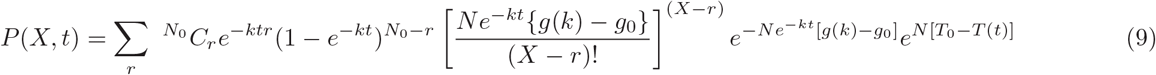

*where, T*(*t*) *is the delay distribution function, such that ∫ ke*^*kt*^ *T*(*t*) *dt* = *g*(*k*) *assuming the rate constants k*_1_ = *k*_2_ = *k*.

***Proof**:* Putting the values of transition probabilities 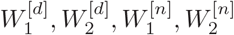 in equation(8), and taking 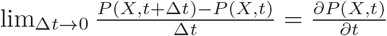, and rearranging the terms we get,

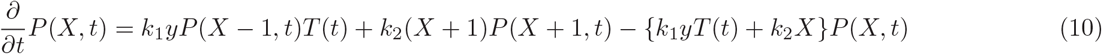

Equation (10) can be solved using generating function method (see *Methods*) [14, 35] to get a general solution of *P*(*X*, *t*). Let the generation function be *G*(*s*, *t*) defined by,

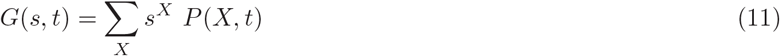

where, *G*(*s*, *t*) satisfy normalization condition, *G*(1, *t*) = 1. Now multiplying equation (10) by *s*^*X*^ and summing over *X* (∑_*X*_), using the definition in equation (11), and rearranging the terms, we get,

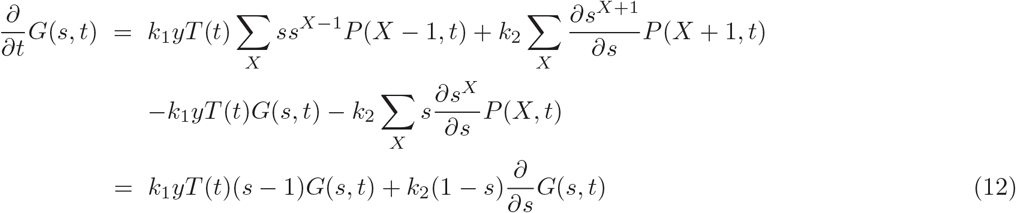

Now, we assume the rate constants to be the same, *k*_1_ = *k*_2_ = *k*, and the variable *X* is translated from the pool of *y* = *N*. Then, we use the transformation, *G*(*s, t*) = *ϕ*(*s*, *t*)*e*^*NsT*(*t*)^, and substituting in equation (12), after rearranging the terms, we have

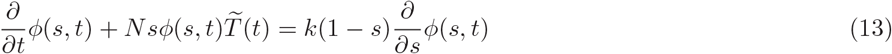

where, 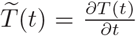. We then use the transformation, (*s* − 1) = *e*^ρ^ and differentiating this with respect to *s* to get, 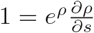. After making the change of variable, *ϕ*(*s*, *t*) → *ψ*(*ρ*, *t*), we have,

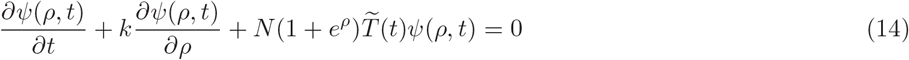

This partial differential equation (14) can be solved using the standard method of characteristics or Lagrange-Charpit equations [36]. The method allows one to define a set of new parameters, *ξ*(*t*, *ρ*) ≡ *ξ*; *η*(*ρ*, *t*) ≡ *η* and *W*(*ξ*, *η*) = *ψ*(*t*(*ξ*, *η*); *ρ*(*ξ*, *η*)), and the following transformations are made from *ψ*(*ρ*, *t*) to *W*(*ξ*, *η*) as: 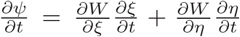; 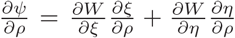. By incorporating these parameters and transformations in (14), the characteristic equation is found to be 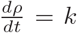. From this, we can get *ρ* = *kt* + *constant*, which allow us to take *η* = *ρ* − *kt*. We can also arrive at *ξ*(*t*, *ρ*) = *ξ* = *t*. The Jacobian for the transformation of variables is non-zero. Finally, substituting these transformations in equation (14), we get the following ordinary differential equation in the new variables,

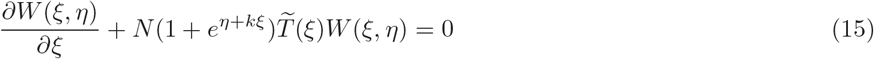

where 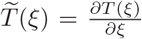. This linear differential equation can be solved using the integrating factor method, where the integrating factor is given by 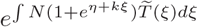. Hence, the general solution of the equation (15) is given by,

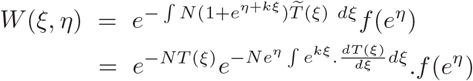

Now integrating by parts and putting *∫ ke*^*kξ*^*T*(*ξ*)*dξ* = *g*(*k*), we get expressions for the mentioned parameters as, 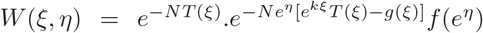; 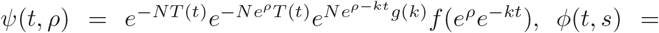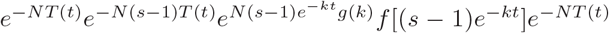. Putting back all these values to the expression for *G*, we have,

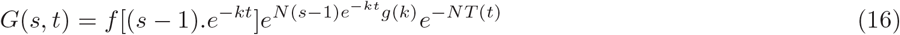

The boundary condition is given by, at *t* = 0, 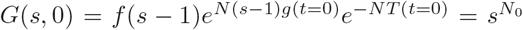. Taking, *q* = *s* − 1, one can get, 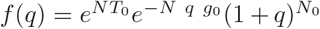, where, *g*(*t* = 0) = *g*_0_ and *T*(*t* = 0) = *T*_0_. Now, after rearrangement of the terms, equation (16) thus becomes,

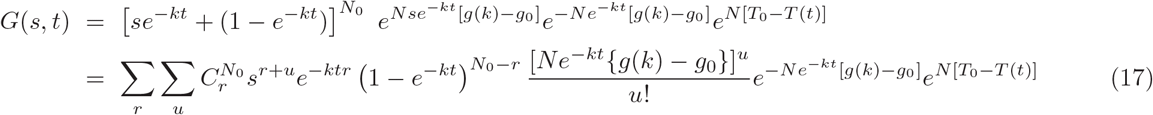

Now, putting *r* + *u* = *X* to equation (17), and using the equation (11), after which equating the coefficients of *s*^*X*^ from both the sides, we arrive at expression for *P*(*X*, *t*),

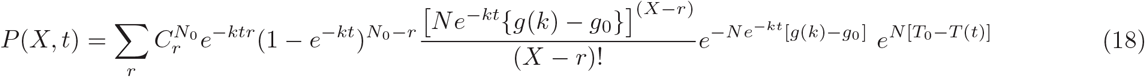

Hence, equation (9) is proven.

#### Theorem 2

*The probability distribution P*(*X*, *t*), *which is the solution of delay master equation of gene regulation, follows Poisson distribution for large N*_0_ *and X given by*,

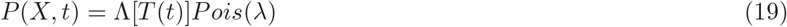

*where*, 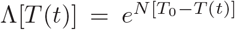 *and* 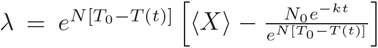 *is the modified mean of the stochastic gene regulatory process.*

***Proof**:* Taking *e*^−*kt*^ = *p* < 1 for significantly large *t*, and then applying *N*_0_ → *large*, the factor in equation (9) can be taken as, 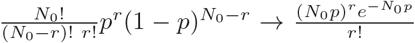, because 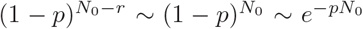 and 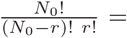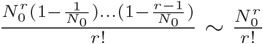. Since, 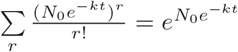, we have for *N*_0_ → ∞ and for large *X*, we have the following *P*(*X*, *t*) given by

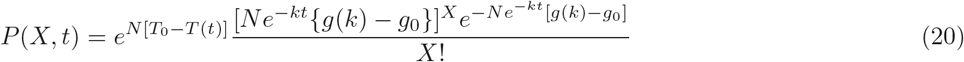

Now, mean ⟨*X*⟩ of the probability distribution *P*(*X*, *t*) can be found from the generating function *G*(*s*, *t*) given in equation (17) which can be written as,

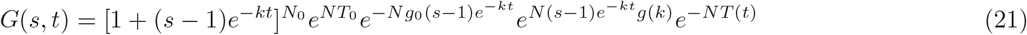

This ⟨*X*⟩ will be given by 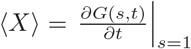. Then using equation (21), we obtain the mean as,

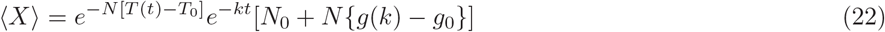

Now, from the equation (20), we can define *λ* = *Ne*^−*kt*^[*g*(*k*) − *g*_0_], and using equation (22), one can write 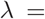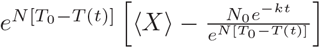. Then upon substitution, the equation (20) becomes,

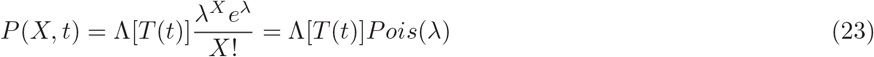

where, 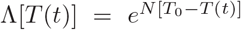. Here, *λ* is the modified mean of the process which depends on the delay time distribution *T*(*t*). Further, as *t* → 0, *λ* → 0, *P*(*X*, *t*) → 0.

#### Lemma 1

*The Fano factor for stochastic gene regulation model is delay time dependent and is given by*,

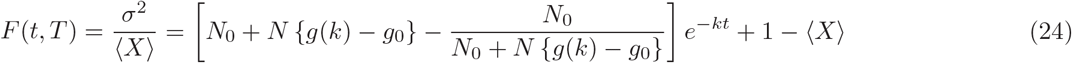

*The condition for sub-Poissonian stochastic process is*,

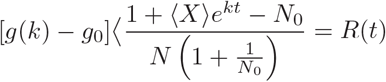

***Proof**:* The variance of the distribution *P*(*X*, *t*) can be calculated directly from the generating function given by equation (21) using the equation,

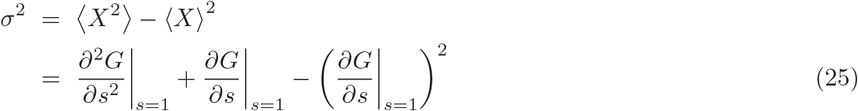

Now, differentiating *G*(*s*, *t*) in equation (21) two times with respect to *s*, and then we put *s* = 1. After rearranging the terms, we obtain,

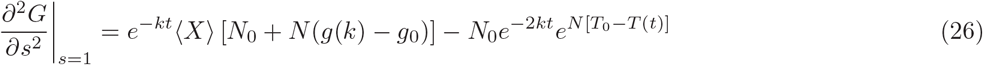

We then put the result in equations (22) and (26) to equation (25) and rearranging the terms we arrive at the following expression for Fano factor [37],

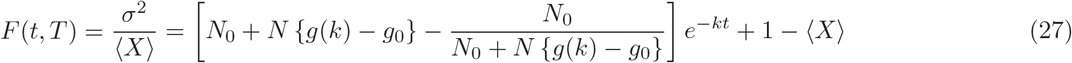

The Fano factor *F* (≥ 0), which is a measure of noise to signal ratio and a dimensionless quantity [37], is similar to coefficient of variation of the probability distribution *P*(*X*, *t*) of the stochastic process. For Poisson process *F* = 1 (absence of correlation of the signals in the process) [37]. Equation (27) shows that delay time has significant role in regulating noise in the stochastic gene expression process. However, existence of deviation of *F* from one in the process indicates the possibility of establishing correlation among the signals in the process: (a) if *F*⟨1, it reveals the suppression of noise such that the process becomes sub-Poissonian process [38]; whereas, (b) if *F*⟩1, it indicates the enhancement of noise in the system [39]. In the stationary case (asymptotic value at *t* → ∞) allows the Fano factor to depend only on delay time *T* associated with *g*(*k*) = *k ∫ e*^*kt*^*T*(*t*)*dt*, and takes the form, *F* → 1, which shows that the system is in Poissonian process for *t* → ∞. Further, if we take the other asymptotic value in *t* (i.e., *t* → 0), we get *F* → 0. Now, the condition for sub-Poissonian stochastic process can be obtained from equation (27), and is,

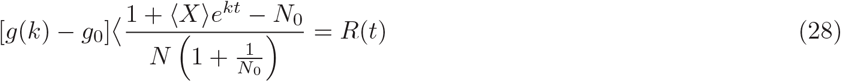

And for noise enhancement process, [*g*(*k*) − *g*_0_]⟩*R*(*t*).

#### Lemma 2

*The probability distribution obtained from delay master equation of gene regulation for constant delay time distribution, T*(*t*) = *T is given by*

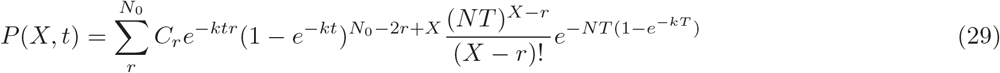

*The Fano factor for this stochastic gene regulation model is given by*,

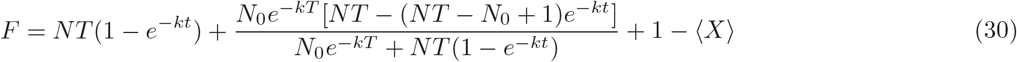

*where, the stochastic gene regulation process follows exact Poisson process at the stationary condition (t* → ∞*), i.e.* lim_*t*→∞_ *F* → 1. *The condition for imparting correlation in the process could be*,

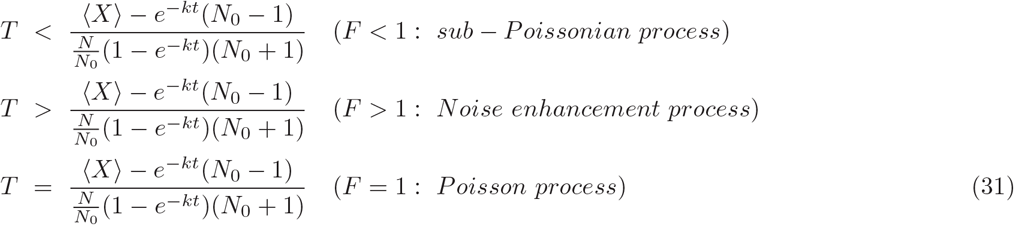

***Proof**:* We know for constant time delay, 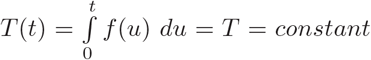. The rate constants are assumed to be equal, *k*_1_ = *k*_2_ = *k* and we have taken *y* = *N*_0_. Now the equation (12) in this case becomes,

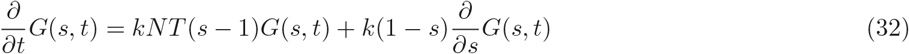

Then we use the transformation *G*(*s*, *t*) = *ϕ*(*s*, *t*)*e*^*Ns*^. Differentiating *ϕ*(*s*, *t*) with respect to *t* and *s* and replacing every terms in equation (32) in functions of *ϕ*, we get the following differential equation in *ϕ*,

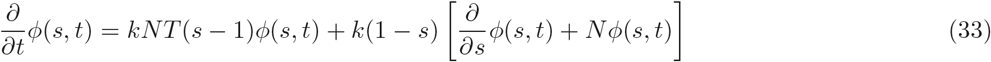

Now, we use the transformation, (*s* − 1) = *e*^*ρ*^ and after differentiating this with respect to *s* we get 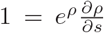. Then, we consider the change of variable, *ϕ*(*s*, *t*) → *ψ*(*ρ*, *t*). Proceeding the change of variable *s* and function *ϕ*(*s*) to corresponding variable *ρ* and function *ψ*(*ρ*) respectively, the following equation can be obtained from equation (33),

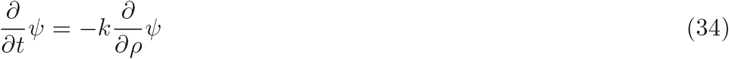

Then, the auxiliary equation of this equation (34) can be written as (*∂ρ* − *k∂t*) = 0, such that, (*ρ* − *kt*) = *constant*. Now, the solution of the equation (34) is given by *ψ*(*ρ*, *t*) = *f*(*e*^*ρ*−*kt*^) = *f*(*e*^**ρ**^.*e*^−*kt*^). Then, after transforming back *ψ*(*ρ*, *t*) → *ϕ*(*s*, *t*) → *G*(*s*, *t*), and using initial condition, 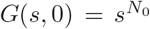, the solution of the equation (34) becomes, *G*(*s*, *t*) = *f* [(*s* − 1)*e*^−*kt*^]*e*^*NT*(*s*−1)^. From this generating function, applying the boundary condition, which is at *t* = 0, 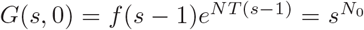, and defining *q* = *s* − 1, we get the functional form, 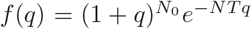 from which we can able to arrive at, 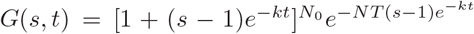. Now, this generating function, therefore, becomes after a few mathematical steps,

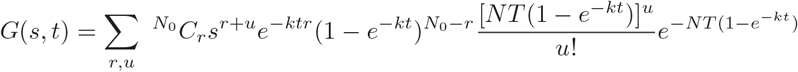

Taking *r* + *u* = *X* such that *u* = *X* − *r*, and then equating coefficients of *s*^*X*^, we get the expression for probability distribution *P*(*X*, *t*),

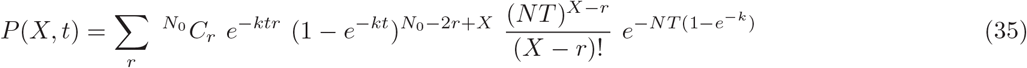

For constant time delay, the observables Mean and Variance can be calculated from the generating function, 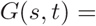 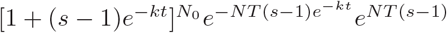 as given below,

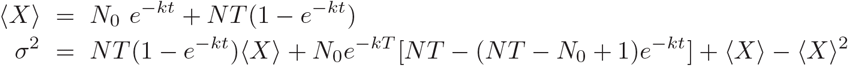

Now, the Fano factor, for this case, can be obtained from these expressions of ⟨*X*⟩ and *σ*, which are given by,

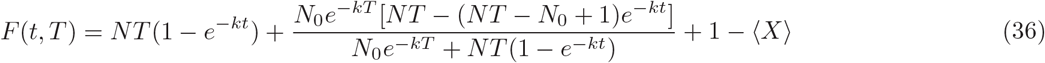

For the stationary case, we can get lim_*t*→−∞_⟨*X*⟩ → *NT*. Then from equation (36), we get, *lim*_*t*→−∞_*F* → 1, which shows that the process is Poisson process. Further, we can also get, lim_*t*→0_ *F* → 0. Now, if the stochastic process is allowed to impart correlation in its process, then *F* < 1 or *F* > 1. Hence, the condition for imparting correlation to the process can be obtained from equation (36) after rearranging the terms,

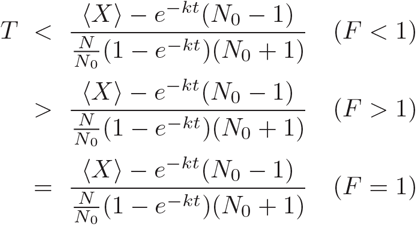

#### Lemma 3

*If the delay distribution obeys linear property in time (i.e., T* = *ct) where, c is a constant and T*(*t* = 0) = 0, *then the probability distribution of the stochastic gene regulation model is given by*,

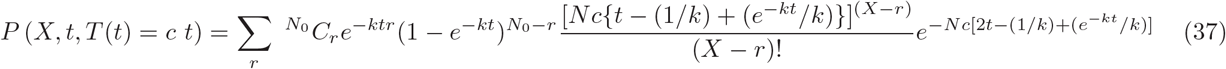

*The Fano factor F in this case can be defined as*,

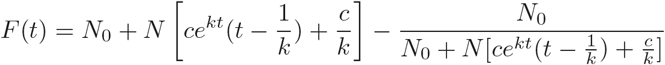

*The condition for introducing correlation in the stochastic process is given by*,

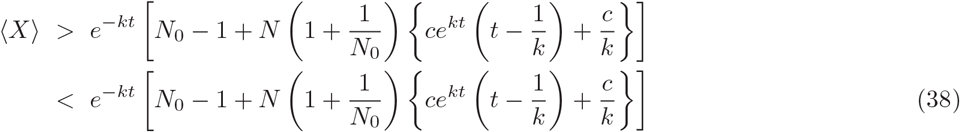

***Proof**:* The function *g*(*k*) for time delay which obeys *T* = *ct* can be calculated as, *g*(*k*) = ∫ *ke*^*kt*^*ctdt* = *ce*^*kt*^(*t* − 1*/k*). The initial condition is given by, *g*_0_ = *g*(0) = −*c/k*. Putting these to the equation (9), we have,

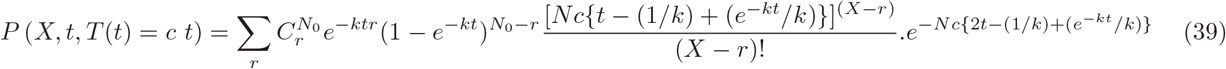

The mean and variance of the gene regulating stochastic process with time delay which obey linear property can be calculated by using equation (21), and putting *T* = *ct*, *g*(*k*) = *ce*^*kt*^(*t* − 1/*k*), and *g*_0_ = −*c/k*. The results are given by,

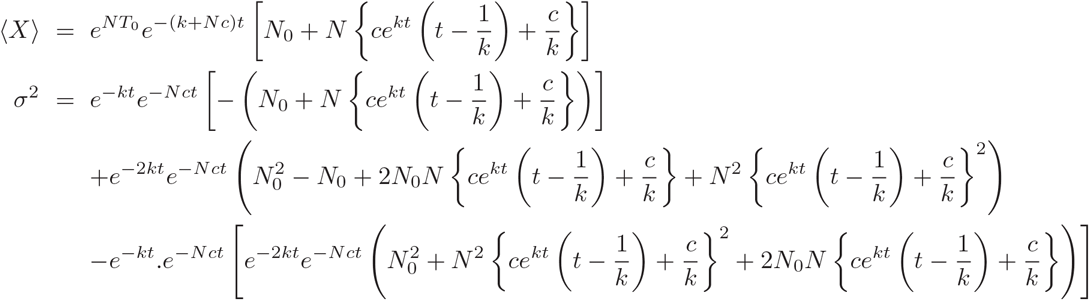

Now, the Fano factor for the process is given by,

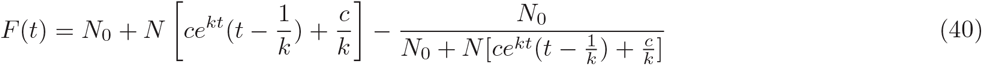

Now if correlation in the stochastic process is associated, then *F* < 1 or *F* > 1. Then from equation (40), we have the conditions for achieving correlation in the process,

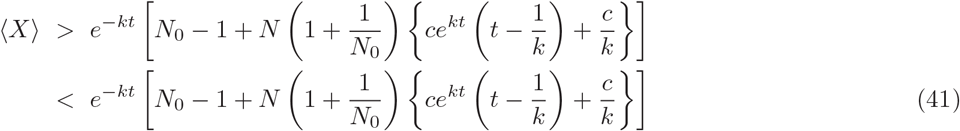

#### Lemma 4

*If the delay distribution is governed by T*(*t*) = Γ*t*^*n*^, *where*, Γ *is a constant and n is an integer with initial condition, T*(*t* = 0) = 0, *then the probability distribution of stochastic Gene Regulatory Model is given by*,

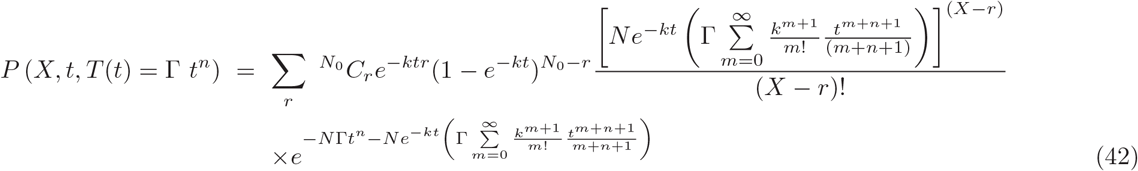

*The Fano factor of this stochastic process is given by*,

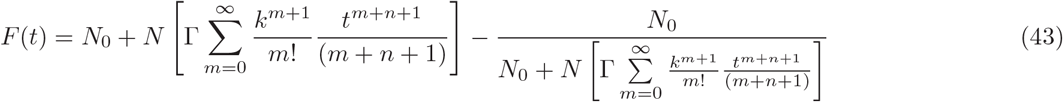

*The condition for allowing to associate correlation in the stochastic process are given by*,

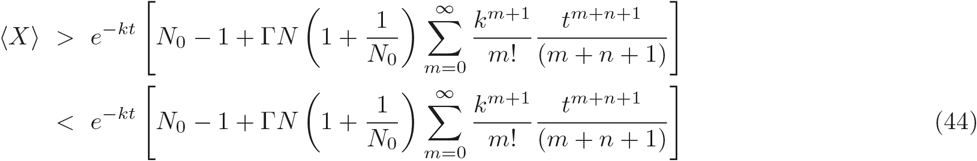

***Proof**:* For the stochastic gene regulatory process, which obey the delay distribution behavior *T* = Γ*t^n^*, we have, 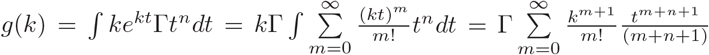. And initial condition is, *g*(0) = 0. Substituting these expressions to equation (9), we prove that the probability distribution is given by equation (42). Now mean and variance can be calculated by substituting the expressions of *g* and *g*_0_ to the equations (21). The results are given by,

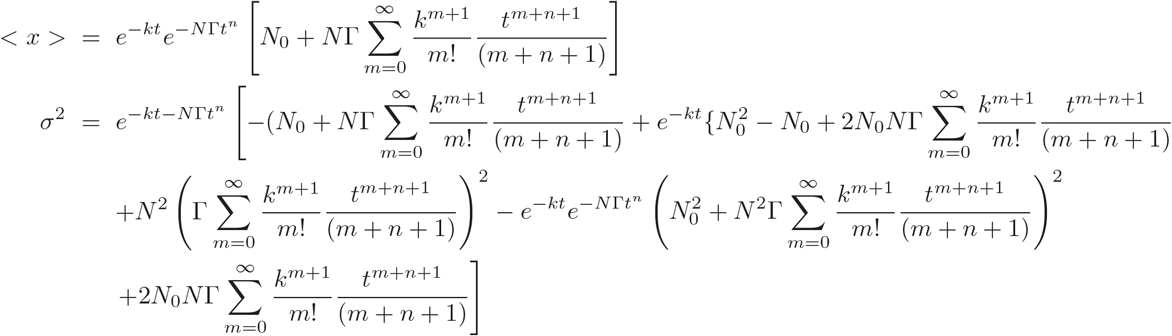

Now, the Fano factor can be calculated from these observables using, 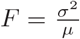 and after rearrangement of the terms, one can prove equation (43). Further, applying the condition for associating correlation in the stochastic gene regulation process, *F* > 1 and *F* < 1, one can prove the equation (44).

### 3.2 Delay induced switching-on oscillations in Gene Regulatory Model

We have taken the Gene Regulatory Model (Table 1) and have used DSSA to study the delay induced behavior of the system variables. The starting of a transcription process is done by binding the promoter region of a gene by *RNA* polymerase. In the elongation phase as the *RNA* polymerase leaves the promoter region, a next *RNA* polymerase can bind to the gene. Since *RNA* polymerase is relatively large, we can depict gene transcription as a non-consuming reaction. The other reaction showing the decay of *mRNA* is taken as the non-delay which is consuming reaction in our simulation using DSSA.

We start our simulation at time t=0.0 (time unit) and simulate till time t=400, and take time delay to be *τ*_*d*_ = 0.0 (without time delay) with initial values and rate constant values listed in Table 1 and 2. We found that the population of polymerase (*P*) remains constant as a function of time, however, the population of *mRNA*(*R*) increases monotonically with respect to time (till *t* ~ 75), and then attained nearly stationary state with fluctuation with very small amplitude as can be seen in Fig. 1-(a). Its corresponding behavior in (*P* − *M*)-plane indicates fixed point nature (Fig. 1 lower left panel). Now, we introduce delay *τ*_*d*_ in the system at fixed initial conditions and same parameter values as listed in Table 1 and 2 by increasing *τ*_*d*_ until we get onset of oscillations in the variables of the system. For small time delays *τ*_*d*_ < 10 we found almost the same behavior of stationary state in the populations of *mRNA*(*R*) and polymerase (*P*) with respect to time as we found in the case of without time delay for this model. As soon as we reach *τ*_*d*_ = 20, suddenly the onset of oscillating behavior starts exhibited in both the dynamics of the variables *P* and *M*, as we can notice in Fig. 1-(a), with well defined varying amplitudes (*A*_*P*_, *A*_*M*_) and time periods of oscillations (*T*_*P*_, *T*_*M*_). As we increase time delay values (*τ*_*d*_ = 30, 40, the oscillatory behavior in the dynamics of *P* and *R* become prominent, (but *A*_*P*_ < *A*_*M*_ and *T*_*P*_ ~ *T*_*M*_) Fig.1-(a). This onset of oscillations both in *P* and *M* dynamics can be seen as broaden limit cycle due to stochastic fluctuations as *τ*_*d*_ → *large* in the (*P* − *M*)-plane (Fig.1-(b)), and could correspond to the activation of transcription process triggered by time delay to enhance gene regulating process. We, then, have plotted amplitude (*A*_*P*_) and time period (*T*_*P*_) with respect to time delay *τ*_*d*_ from where they start to oscillate (Fig.1-(c) and (d)), with error bars, and show increase in both amplitude and time period of *P* as time delay *τ*_*d*_ increases. This results are supported by experimental observation on the onset of oscillation in cyclic expression of mRNA and protein of serum treated cultured cells Hes1, a basic helix-loop-helix factor, with a certain periodicity [43].

**FIG. 1:**
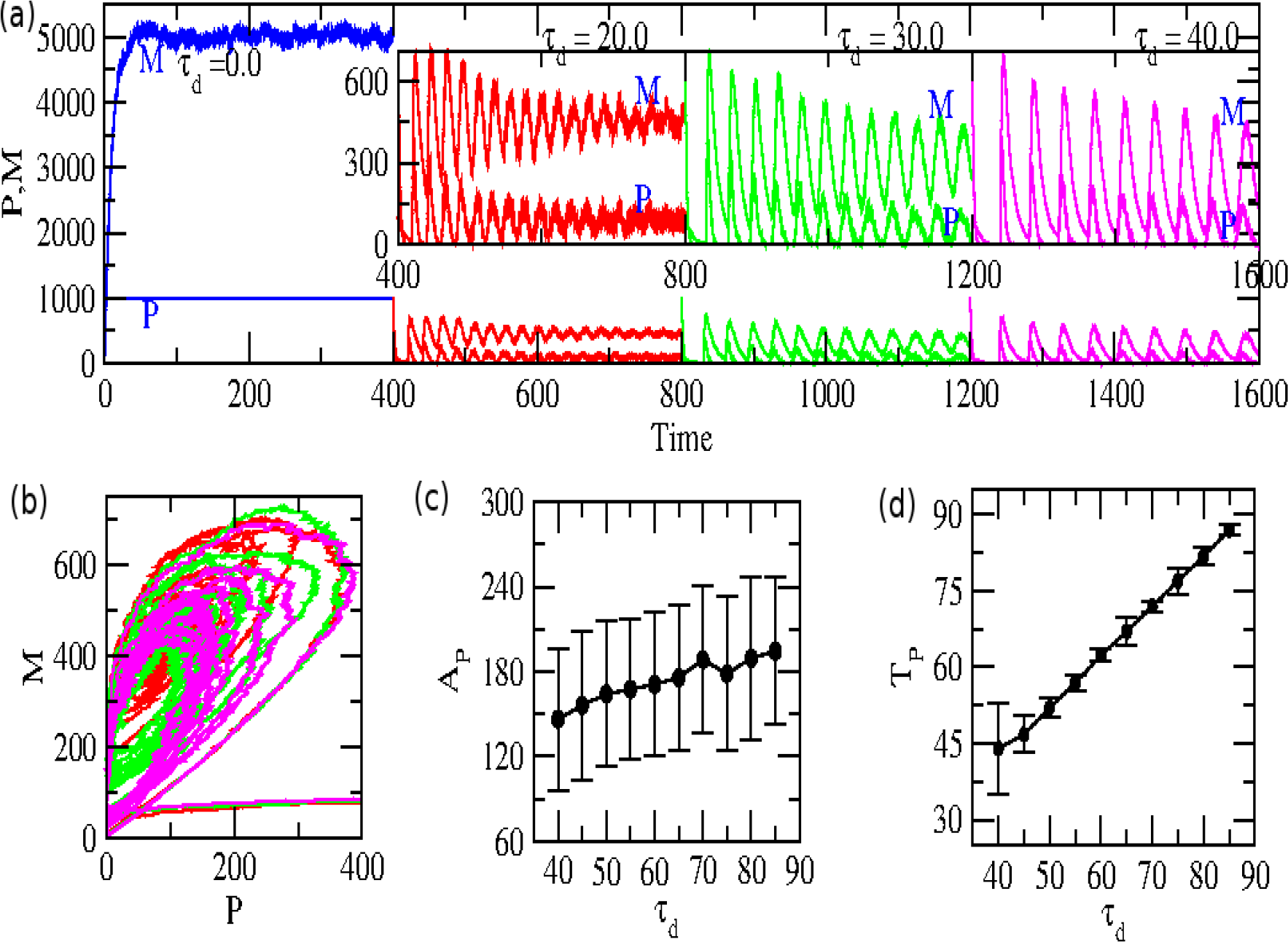
*Delay induced switching-on oscillations in gene regulatory model: Results of delay stochastic simulation of gene regulatory model with multiple time delay values. Initial population of of the variables and parameter values are P=1000, M=100, k*_1_ = 0.5*t*^−1^, *k*_2_ = 0.1*t*^−1^. *Duration of simulated time = 400.0. The values of time delay taken are: (a) τ_d_* = 0.0 *(blue)*, *τ_d_* = 20.0*t (red), τ_d_* = 30.0*t (green) and τ_d_* = 40.0*t (pink), (b) Two dimensional plot of populations of M with P, (c) Amplitude (A) of P versus τ*_*d*_ *with error bars, (d) Time period (T) of P versus τ*_*d*_ *with error bars.*

### 3.3 Delay induced optimization of cellular rhythm

We have taken reduced three variables circadian rhythm model regulated by three clock proteins, namely, cytosolic clock protein (*P*_*C*_), nucleus clock protein (*P*_*N*_), and mRNA (*M*) [40]. In this simple model, the clock protein (*P*_*C*_) enters the nucleus after getting signal from the clock regulatory mechanism at cytosol to regulate nuclear clock protein (*P*_*N*_) and to repress gene into mRNA (*M*). This regulatory clock mechanism is based on negative feedback loop developed by a clock protein (*P*) on its gene expression [40]. The detailed list of biochemical reactions, associated parameters, and their values used in our simulations are provided in Table 1 and 2. We took the sixth reaction (Table 1) as the delay reaction. Using these six reactions, and following the same procedure we did in the case of gene regulation, we can construct Master equation from detailed balance condition,

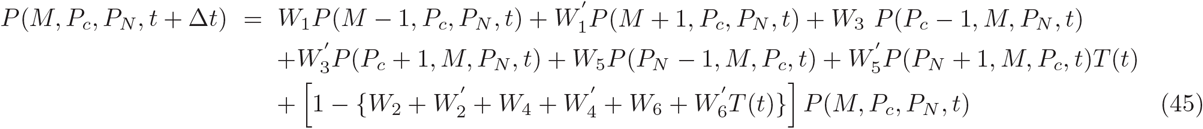

where, {*W*} are the transition probabilities of the state vector, *X* = [*M*, *P*_*C*_, *P*_*N*_]^−1^. Now putting the expressions for {*W*} from the reaction network, we arrive at the following Master equation,

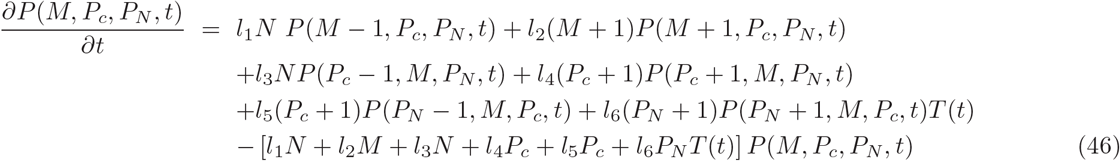

Now we use DSSA to solve this DSME numerically by simulating the corresponding reactions which construct the above DSME to study the impact of delay *τ*_*d*_ to the system dynamics. We consider a set of parameter values as listed in Table 1 and 2 which exhibit stationary behavior at *τ*_*d*_ = 0 during the time interval [0, 400]. We found that till *τ*_*d*_ ≤ 2, the dynamics of *P*_*C*_, *P*_*N*_, *M* exhibit stationary behavior as shown in Fig.2 -(a). However, if time delay is *τ*_*d*_ ≥ 3, the system’s variables *P*_*C*_, *P*_*N*_, *M* show significant oscillations with well defined amplitude and time period (Fig.2 -(a)). It is also observed that the amplitudes and time periods of oscillations of these system’s variables increase as *τ*_*d*_ increases. The exhibition of fixed point (corresponding to *τ* ≤ 2) behavior to broaden sustain oscillations as *τ*_*d*_ increases shows the evidence of switching on of oscillations to the system driven by time delay. Further, the results also indicate that *τ*_*d*_ has to optimize such that the time periods of clock genes (*P*_*C*_, *P*_*N*_, *M*) be maintained properly ~ 24*hours* time period for proper regulation of the clock mechanism. These results of emergence of oscillation in the clock gene is supported by the experimental observation of oscillations in CPer2 clock gene of chicken pineal when phase delay is introduced to transcription factor E4bp4, and then light-dependent suppression of the rhythm [44]. Further, there are many experiments done on mammalian circadian rhythm which have shown the onset of oscillation induced by time delay [45], specially in *period 3* clock protein [46]. From these results, it seems that natural mechanism of optimization of time period of oscillation and amplitude of the clock proteins is inbuilt in all living organisms for proper functionality of the system, otherwise it could lead to various diseases. This onset of oscillation with proper amplitude and time period in the clock genes is an important factor, and can be maintained by time delay *τ*_*d*_ without affecting the regulatory network. It is also quite possible that delay could be an inbuilt mechanism to maintain the system’s properties and behavior when it is externally perturbed.

**FIG. 2:**
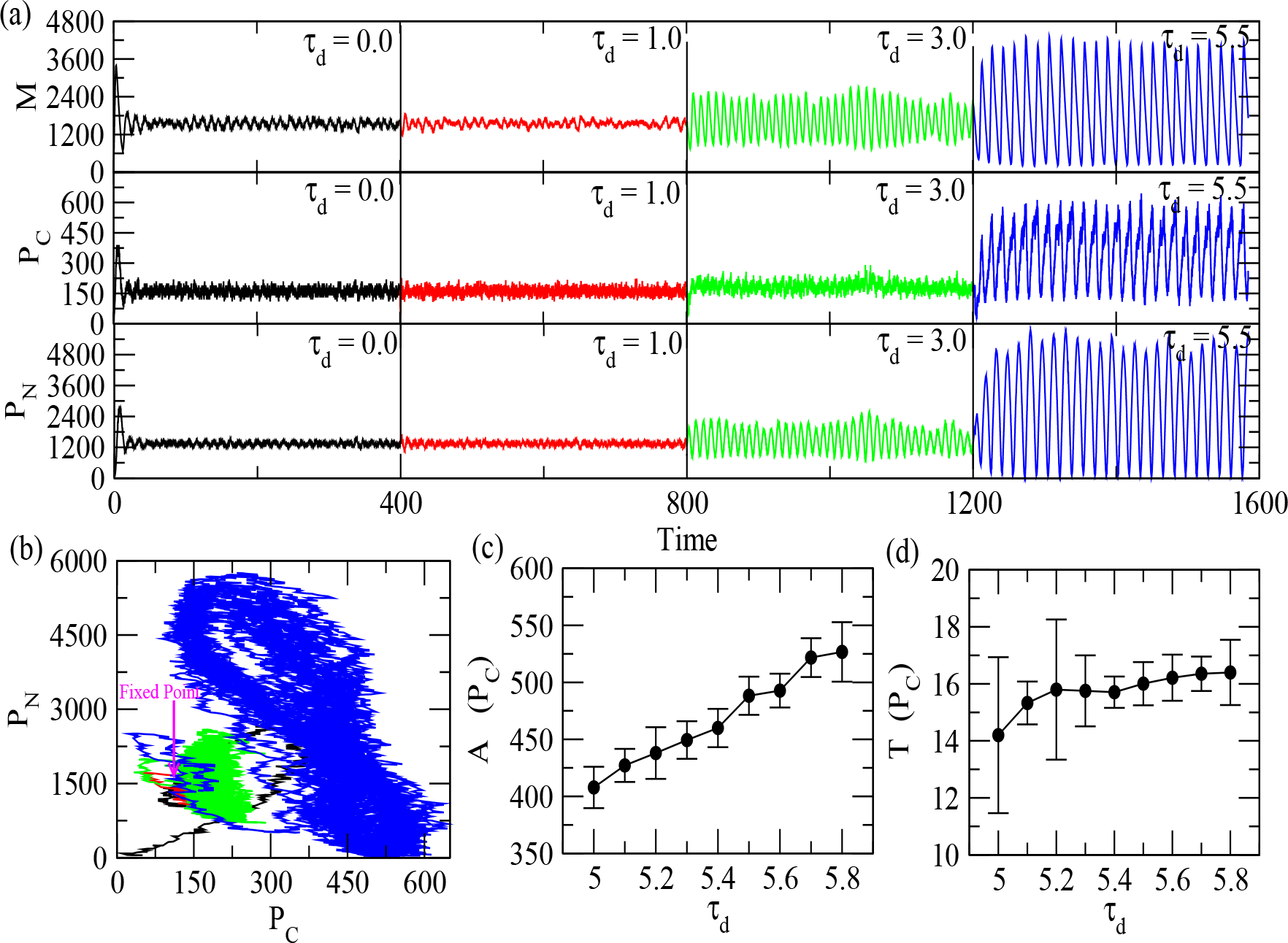
*Delay induced onset of oscillations in circadian rhythm model: Initial populations of the molecular species in the model and parameter values, M* = 50, *P*_*C*_ = 50, *P*_*N*_ = 50, *v*_*s*_ = 1.6*nMh*^−1^, *v*_*m*_ = 0.505*nMh*^−1^, Ω = 1000.0, *K*_1_ = 1.0*nM, n* = 4, *K*_*m*_ = 0.5*nM, k*_*s*_ = 0.5*h*^−1^, *u_d_* = 1.4*nMh*^−1^, *K_d_* = 0.13*nM, k*_2_ = 0.6*nMh*^−1^, *k*_3_ = 5.0*nMh*^−1^, *(a) Time intervals are [0.0,400.0], [400.0,800.0], [800.0,1200.0] and [1200.0,1600.0]. Corresponding time delays are τ*_*d*_ = 0.0, *τ*_*d*_ = 1.0, *τ*_*d*_ = 3, *and τ*_*d*_ = 5.5, *(b) Two dimensional plot of population of P*_*C*_ *with population of P*_*N*_, *(c) Plots of amplitude (A) of PC with respect to τ*_*d*_ *with error bars, (d) Plot of time period (T) of PC as a function of τ*_*d*_ *with error bars.*

### 3.4 Delay induced onset of activation of repressilator dynamics

The repressilator model involves three genes. The inhibition among the genes occurs in a circular manner; the protein of first gene inhibits the second gene, new protein from the second gene inhibits the third gene, and finally protein of this gene inhibits the first gene [24]. The model consists of six reactions as listed in Table 1, and detail parameters and their values used in our simulation are provided in Table 1 and 2. We considered reactions (1), (3) and (5) in the Table 1 for this model to be the delay reactions, and the rest three reactions as non-delay reactions. Following the same procedure described in *Methods*, the DSME of the symmetrical repressilator model is given by,

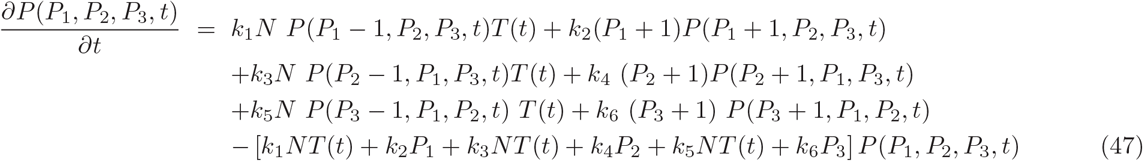

We now use DSSA (see *Methods*) to simulate the model by using the reactions which correspond to this Master equation (Table 1 and 2). Parameter *a* is constant and parameter *b* is cooperativity and reflects multimerization of the protein required to affect the promoter. We have done the simulation with initial population *P*_1_ = *P*_2_ = *P*_3_ = 1. The simulation is done for a duration of 1600 time steps by introducing different time delay *τ*_*d*_ values at various time intervals of duration 400 each (Fig:3). For time interval [0, 400], we took time delay *τ*_*d*_ = 0.0 (non-delay case) for the simulation. The results show a random fluctuations in the species variables (*P*_1_, *P*_2_, *P*_3_) with significantly very small random amplitudes and time periods with respect to time. Then for the next time interval [400, 800], we introduced time delay *τ*_*d*_ = 1.0 with the same parameter values as done in the case of non-delay case. We found the emergence of weak oscillatory behavior (oscillation with early defined amplitudes and time periods) in variables’ dynamics with significantly bigger amplitude and time period but random fluctuations in the populations of the variables with respect to time (see Fig.3-a). During the next time interval [800, 1200], the value of time delay is increased *τ*_*d*_ = 5.0 with the same parameter values as before. The results now show that the emerged oscillations in the variable dynamics become prominent with well defined amplitudes and time periods in both *P*_1_ and *P*_2_ with significantly reduced fluctuations in their amplitudes and time periods (Fig:3-(a)). Now, for the next time interval [1200, 1600] we increased the time delay *τ*_*d*_ = 8.0, and we found that the oscillation is well defined with prominent amplitudes and time periods in the dynamics of *P*_1_ and *P*_2_. The behavior of *P*_1_ and *P*_2_ in the (*P*_1_ − *P*_2_)-plane show fixed point behavior when *τ*_*d*_ = 0, and then trigger to broaden limit cycle behavior as *τ*_*d*_ → *large* (Fig.3-(b)). Since the behavior of the three variables (*P*_1_, *P*_2_, *P*_3_) are similar, we present the behavior of amplitude 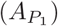 and time period 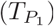 with respect to time delay *τ*_*d*_ (Fig.3 (c) and (d)), with error bars. The behavior of 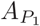 becomes stationary as *τ*_*d*_ → *large <* 13 indicating onset of limit cycle driven by *τ_d_*. However, 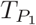 increases as *τ*_*d*_ increases. These observation of inducing oscillation by repressilator model is supported by experimental findings of oscillatory behavior in three clock gene repressilator circuit in Arabidopsis [47].

**FIG. 3:**
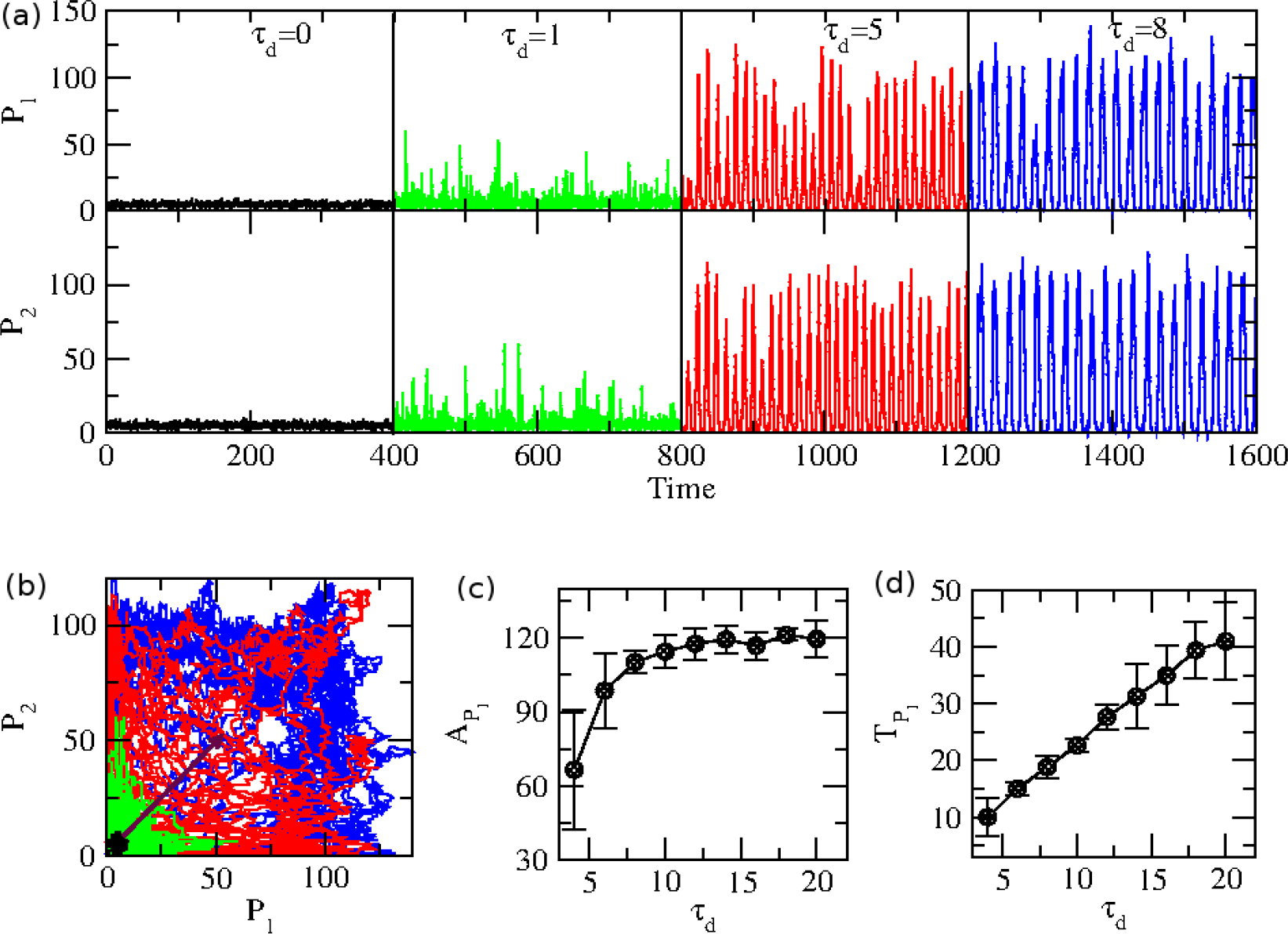
*Delay induced activation of states in repressilator model. The initial populations of the clock variables and parameter values are, P*_1_ = *P*_2_ = *P*_3_ = 1, *a=100.0, b=2.0, k*_2_ = 1.0*t*^−1^, *k*_4_ = 1.0*t*^−1^, *k*_6_ = 30.0*t*^−1^. *(a) Plots of P*_1_ *and P*_2_ *as a function of time with different time intervals for various τ*_*d*_, *[0.0,400.0], [400.0,800.0], [800.0,1200.0] and [1200.0,1600.0] with corresponding time delays are τ*_*d*_ = 0.0, *τ*_*d*_ = 1.0, *τ*_*d*_ = 5.0, *and τ*_*d*_ = 8.0, *(b) Two dimensional plots of P*_1_ *versus P*_2_, *(c) Plots of amplitude (A) of P*_1_ *and, (d) time period (T) of P*_1_ *as a function of τ*_*d*_ *with error bars.*

The results clearly show that the introduction of time delay may lead to the onset of cooperative nature among the behavior of the three genes, which may be due to delay induced coherent information processing reflected in the synchronized behavior of the prominently oscillating dynamics of the genes. The delay induced switching on of the oscillations in the system’s variables (*P*_1_, *P*_2_, *P*_3_) might trigger the system to active state, opens up possibilities of accessing information among the genes (even though they are inhibiting each other in symmetric circular way) because of the oscillating nature exhibited, and can establish coherence among them with significantly better information processing.

### 3.5 Delay induced oscillation death in chemical oscillator

We used brusselator model, which is a chemical oscillator developed by Ilya Prigogine and consists of four reaction channels (Table 1), for our study [41]. This model could generate stable oscillations in the dynamics of species variables [16]. It is similar to auto-catalytic reaction model whose dynamics in the system variables and reaction mechanisms are similar to that of auto-catalytic reactions [42]. This chemical oscillator is a two variable model, namely *Y*_1_ and *Y*_2_. Following the same procedure described in the *Methods*, we can construct DSME from the four reactions listed in Table 1, where, we introduce delay in the third reaction, and the remaining three reactions are taken as non-delay reactions. The DSME of brusselator model is given by,

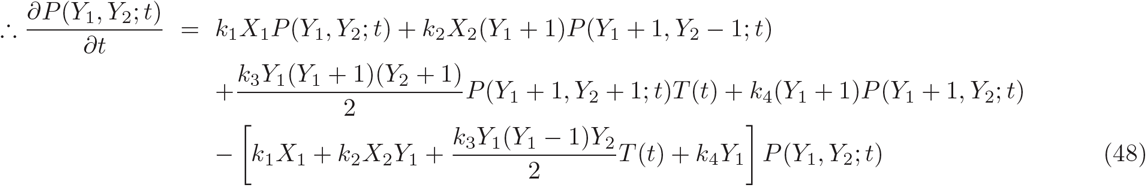

We have simulated this model by taking the reaction channels from which the DSME was constructed taking initial population of *Y*_1_ = 1000, *Y*_2_ = 2000, *X*_1_ = *Y*_1_, and *X*_2_ = *Y*_2_. Other parameter values for the simulation of the system are provided in Table 1 and 2. The results of time delay *τ*_*d*_ = 0.0 shows well defined stochastic oscillation in the dynamics of *Y*_1_ and *Y*_2_ (see Fig.4(a) and (e)). When the induced delay is very small (*τ*_*d*_ = 0.005, 0.5, 1 *<* 2.5 we could see the oscillating behavior in the dynamics of system parameters. However, if introduced time delay is large *τ* = 2.5, the exhibited oscillation in both the dynamics of *Y*_1_ and *Y*_2_ become vanished leaving few random spikes and rest with random behavior with time (Fig.4(b) and (f)). With further increase in the value of *τ*_*d*_ ≥ 2.5, similar switching-off of oscillations nature exhibited in both *Y*_1_ and *Y*_2_ dynamics (Fig.4(c) and (g)). As *τ*_*d*_ → *large*, the oscillation completely vanished with few spikes, and left only random behavior(Fig.4(d) and (h)). This could be the state of the system where system is in stationary state.

**FIG. 4:**
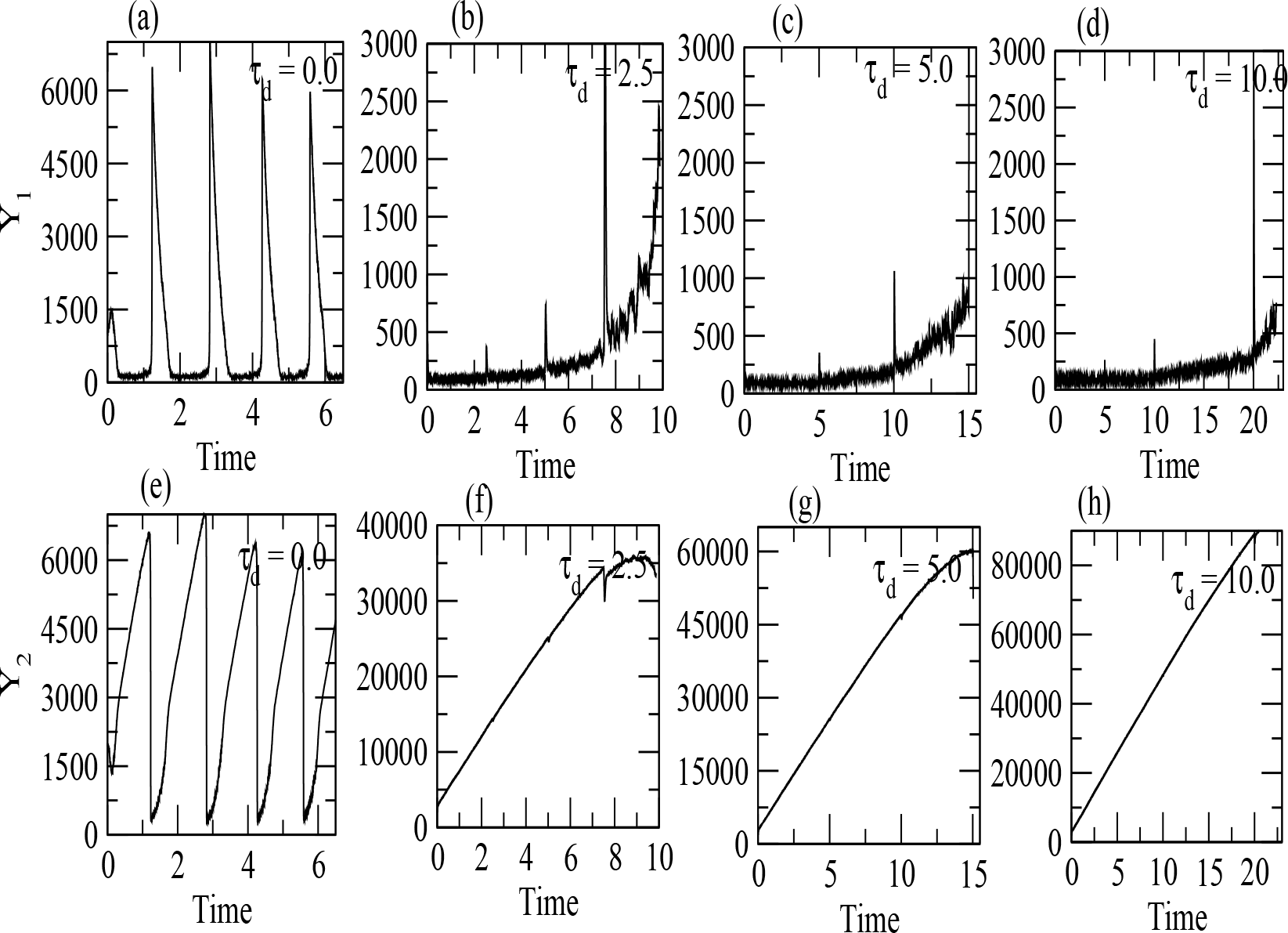
*Delay induced switching-off oscillations in Brusselator: The initial populations and parameter values taken are: Y*_1_ = *1000*, *Y*_2_ = *2000*, *X*_1_ = *Y*_1_, *X*_2_ = *Y*_2_, *a=100.0, b=2.0, k*_1_ = 5.0*t*^−1^, *k*_2_ = 0.025*t*^−1^, *k*_3_ = 0.00005*t*^−1^, *k*_4_ = 5.0*t*^−1^. *Plots of Y*_1_ *population for different time delay (a) τ*_*d*_ = 0.0, *(b) τ*_*d*_ = 2.5 *(c), τ*_*d*_ = 5.0, *(d) τ*_*d*_ = 10.0. *Similarly plots of Y*_2_ *population for different time delay values (e) τ*_*d*_ = 0.0, *(f) τ*_*d*_ = 2.5 *(g) τ*_*d*_ = 5.0 *and (h) τ*_*d*_ = 10.0.

### 3.6 Delay induced bistable states and switching mechanism

Consider symmetrical repressilator model we discussed in the previous section and described in Table 1 and 2 with all the same parameter values in the simulation using DSSA, but the delay time *τ*_*d*_ ≥ 20. The results for *τ*_*d*_ = 20 show that the random behavior in the dynamics of the system variables *P*_1_, *P*_2_, *P*_3_ at *τ*_*d*_ = 0 (non-delay behavior) becomes two state behavior or bistable states in the dynamics (Fig.5). As *τ*_*d*_ increases, the emergent two states are more prominent and the width of the bistable states becomes larger. However, the population of the system’s variables at the two states do not change significantly even though the dynamics show stochastic fluctuations. This reaction set up of the ring type repressilator is quite mimic with the proposed experimental design in Escherichia coli cells (in the production and decay of mRNA signals) and results obtained in the work by Kim and Winfree [48].

**FIG. 5:**
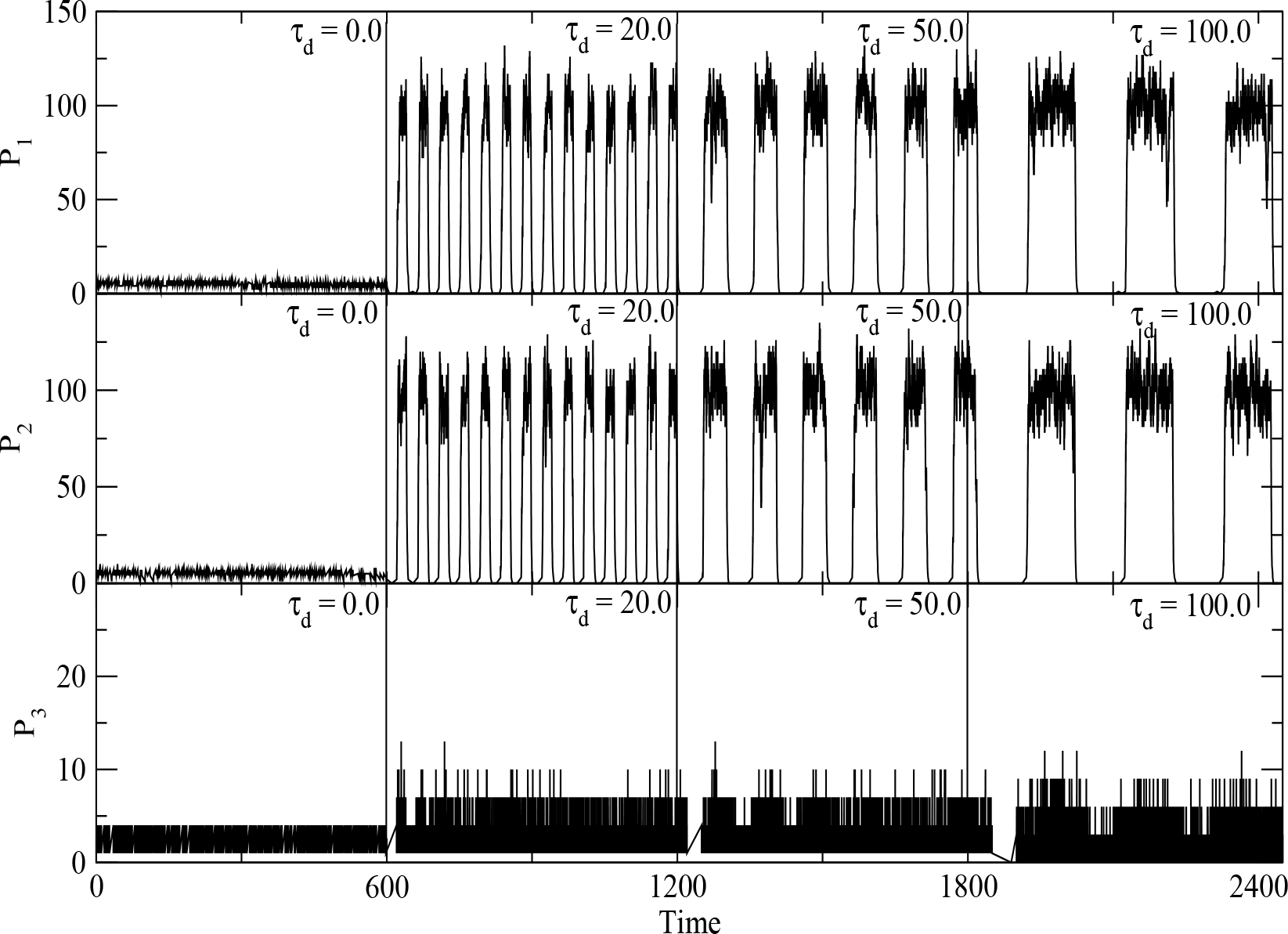
*Delay induced bistable states repressilator model: Initial population of the variables and parameter values are: P*_1_ = *P*_2_ = *P*_3_ = 1, *a=100.0, b=2.0, k*_2_ = 1.0*t*^−1^, *k*_4_ = 1.0*t*^−1^, *k*_*6*_ = 30.0*t*^−1^. *Plots of P*_1_, *P*_2_, *P*_3_ *as a function of time for time intervals [0.0,600.0], [600.0,1200.0], [1200.0,1800.0] and [1800.0,2400.0]. Corresponding values of time delay taken are, τ*_*d*_ = 0.0, *τ*_*d*_ = 20.0, *τ*_*d*_ = 50.0 *and τ*_*d*_ = 100.0.

The role of time delay is indeed quite complex, and system dependent. In this situation, larger time delay induces correlated states of the system’s variables even though the genes are inhibiting among them in a circular fashion. Further, this time delay may create the coherent states to extend for longer time to stay which may probably help in decision making mechanism of the interacting genes. Decision making mechanism is generally facilitated by allowing the system to evolve in bistable states process, and is generally an inbuilt property in most of the biological systems. Introduction of time delay may probably open up the already existed bistable state or created new bistable state for the system for making decision.

## 4 Conclusion

Delay is everywhere how small or big it is. Delayed reactions, which take certain significant time to finish after they are initiated, play an important role in regulating stochastic biological systems. They could control the behavior of the system’s dynamics, and could trigger the system to active state. Since the delayed reactions follow non-Markovian process, the Master equation of any system, which consists of both delay and non-delay reactions, needs to be modified by DSME incorporating time delay parameter in the formalism. This DSME equation is in general difficult to solve analytically except for simple ones. We considered Gene Regulatory Model for which DSME can be solved analytically, and showed that for stationary process the system follows Poisson like process which is one universal class of stochastic process. Then we also can get various conditions on time delay parameter for which one can trigger various mechanisms, such as, coherence, correlated activity in the process etc. We can also understand the possibility of noise induced control of the system driven by time delay. For systems which can not be solved analytically, one needs to use delayed stochastic simulation algorithm proposed by Barrio et al because the stochastic simulation algorithm proposed by Gillespie does not give correct simulation result. The delayed stochastic simulation algorithm now can reveal the importance of delay in the study of complex biological processes significantly at molecular level [24]. We have applied this DSSA to various biological system models and showed various possible roles of time delay.

**Table 1.** List of chemical channels and propensity functions

| ModelName | Reactions | Propensity functions |
| --- | --- | --- |
| Gene Regula-<br>tory Model | 1- $P \xrightarrow{k_1} M + P$<br>2- $M \xrightarrow{k_2} \phi$ | 1- $k_1 P$<br>2- $k_2 M$ |
| Circadian<br>Oscillation<br>Model | 1- $\phi \xrightarrow{l_1} M$<br>2- $M \xrightarrow{l_2} \phi$<br>3- $\phi \xrightarrow{l_3} P_C$<br>4- $P_C \xrightarrow{l_4} \phi$<br>5- $P_C \xrightarrow{l_5} P_N$<br>6- $P_N \xrightarrow{l_6} P_C$ | 1- $v_s \Omega \frac{(K_1 \Omega)^2}{(v_s \Omega)^2 + (P_N)^n}$<br>2- $v_m \Omega \frac{M}{(K_m \Omega) + M}$<br>3- $k_s M$<br>4- $v_d \Omega \frac{P_C}{(K_d \Omega) + P_C}$<br>5- $k_3 P_C$<br>6- $k_2 P_N$ |
| Symmetrical<br>Repressilator<br>Model | 1- $\phi \xrightarrow{k_1} P_1$<br>2- $P_1 \xrightarrow{k_2} \phi$<br>3- $\phi \xrightarrow{k_3} P_2$<br>4- $P_2 \xrightarrow{k_4} \phi$<br>5- $\phi \xrightarrow{k_5} P_3$<br>6- $P_3 \xrightarrow{k_6} \phi$ | 1- $\frac{a}{1 + (P_3(t)^b)}$<br>2- $k_2 P_1$<br>3- $\frac{a}{1 + (P_1(t)^b)}$<br>4- $k_4 P_1$<br>5- $\frac{a}{1 + (P_2(t)^b)}$<br>6- $k_6 P_1$ |
| Brusselator<br>Model | 1- $X_1 \xrightarrow{k_1} Y_1$<br>2- $X_2 + Y_1 \xrightarrow{k_2} Y_2 + Z_1$<br>3- $2Y_1 + Y_2 \xrightarrow{k_3} 3Y_1$<br>4- $Y_1 \xrightarrow{k_4} Z_2$ | 1- $k_1 * X_1$<br>2- $k_2 * X_2 * Y_1$<br>3- $k_3 * \frac{Y_2 * (Y_1 - 1)}{2}$<br>4- $k_4 * Y_1$ |

**Table 2.** List of parameters and values

| ModelName | Symbols | Values |
| --- | --- | --- |
| Gene Regula-<br>tory Model | $P$ (Polymerase)<br>$M$ (mRNA)<br>$k_1$ (Transcription rate)<br>$k_2$ (Translation rate) | 1000(initial)<br>100(initial)<br>$0.5t^{-1}$<br>$0.1t^{-1}$ |
| Circadian<br>Oscillation<br>Model | $M$ (mRNA)<br>$P_C$ (Cytosolic Clock Protein)<br>$P_N$ (Nucleus Clock Protein)<br>$\Omega$ (System Size)<br>$v_s$ (Maximum transcription rate)<br>$v_m$ (Maximum degradation rate)<br>$k_1$ (Transportation rate constant into nucleus)<br>$k_2$ (Transportation rate constant into cytosol)<br>$k_3$ (Michaelis constant for reversible phosphorylation)<br>$k_m$ (Michaelis constant)<br>$k_d$ (Michaelis constant)<br>$k_s$ (Synthesis rate of protein)<br>$u_d$ (Degradation rate for protein)<br>$n$ (Degree of cooperativity) | 50(initial)<br>50(initial)<br>50(initial)<br>1000<br>$1.6nMh^{-1}$<br>$0.505nMh^{-1}$<br>1.0nM<br>$0.6nMh^{-1}$<br>5.0 nM<br>0.5nM<br>0.13nM<br>$0.5h^{-1}$<br>$1.4nMh^{-1}$<br>4 |
| Symmetrical<br>Repressilator<br>Model | $P_1$ (Protein of first gene)<br>$P_2$ (Protein of second gene)<br>$P_3$ (Protein of third gene)<br>$a$ (Constant)<br>$b$ (Cooperativity)<br>$k_2$ (Translation rate constant for $P_1$)<br>$k_4$ (Translation rate constant for $P_2$)<br>$k_6$ (Translation rate constant for $P_3$) | 1(initial)<br>1(initial)<br>1(initial)<br>100<br>2<br>$1.0t^{-1}$<br>$1.0t^{-1}$<br>$30.0t^{-1}$ |
| Brusselator<br>Model | $X_1$ (Chemical species)<br>$X_2$ (Chemical Species)<br>$Y_1$ (Chemical species)<br>$Y_2$ (Chemical Species)<br>$k_1$ (Reaction rate)<br>$k_2$ (Reaction rate)<br>$k_3$ (Reaction rate)<br>$k_4$ (Reaction rate) | 1000(initial)<br>2000(initial)<br>1000(initial)<br>2000(initial)<br>$5.0t^{-1}$<br>$0.025t^{-1}$<br>$0.00005t^{-1}$<br>$5.0t^{-1}$ |

The role of delay is found to be multi functional and seems to be system dependent. In gene regulating process, introduction of time delay triggers the onset of well defined oscillatory state at sufficiently large values of time delay. These oscillatory states could be the states at which the system is in active state, such that, the efficiency of the performance of the system at these states is optimized. Similar onset of oscillating states driven by time delay are also obtained in circadian rhythm as well as in symmetrical repressilator. From the study of circadian rhythm, introduction of time delay to the reaction system allows the system to optimize the rhythm properties, such as, time period, amplitude etc so that the system performs active and normal. On the other hand, switch on of oscillating states driven by time delay to repressilator allows the system to establish correlated states among the genes, which are inhibiting to each other in circular manner, and the genes comfortably process signal to keep the system active. On the hand, time delay can switch off the oscillation in the case of chemical oscillator, namely, brusselator, which could correspond to the inactive state or system failure.

One of the most important roles of time delay in regulating a system could be the possibility of creating coherent bistable states, where, the system can stay longer at each state for making decisions for the fate of the system. In such situation, decision making mechanism provided by the time delay is quite important, specially when the system is perturbed by external fluctuations or in disease state attacked by pathogens etc. However, it is also important to devise general methods to solve Master equation as well as DSME analytically to understand about the roles of time delay in a more deeper level.

## 5 Acknowledgments

S.N.S. (award no. 09/263(1096)/2016-EMR-I) is financially supported by Council of Scientific and Industrial Research, New Delhi, India. A.L.C. is a DST-Inspire Fellow (IF180043) and acknowledges Department of Science and Technology (DST), Government of India for financial support under Inspire Fellowship scheme (order no:DST/INSPIRE Fellowship/[IF180043]). M.Z.M. financially supported by Department of Health and Research, Ministry of Health and Family Welfare, Government of India under young scientist FTS No. 3146887. R.K.B.S. acknowledges UPE-II, sanction no. 101, India, for providing financial support.

## I. AUTHOR CONTRIBUTIONS

S.N.S., M.Z.M. and R.K.B.S conceived the model. A.L.C. and R.K.B.S. did analytical work. S.N.S. and R.K.B.S. did the numerical experiment and prepared the figures of the numerical results. S.N.S. A.L.C., M.Z.M. and R.K.B.S. analyzed and interpreted the analytical as well as simulation results. All authors wrote and approve the final manuscript.

## II. ADDITIONAL INFORMATION

**Competing financial interests:** The authors declare no competing financial interests.

